# The effects of muscle fibre type distribution on gait biomechanics: A predictive simulation study

**DOI:** 10.64898/2026.04.13.718234

**Authors:** Torstein E. Dæhlin, Stephanie A. Ross, Friedl De Groote, James M. Wakeling

**Affiliations:** Department of Biomedical Physiology & Kinesiology, Simon Fraser University, Burnaby; Faculty of Kinesiology University of Calgary, Calgary; Department of Movement Sciences, KU Leuven, Leuven

**Keywords:** Musculoskeletal modelling, Muscle fibres, Motor coordination, Locomotion

## Abstract

Muscle fibre type distribution influences both metabolic and contractile properties of muscles and its influence on gait biomechanics is difficult to investigate *in vivo*. This study aimed to predict the influence of muscle fibre type distribution on the metabolic cost and biomechanics of simulated gait. A muscle model that independently represented slow and fast twitch fibres was implemented in a predictive musculoskeletal simulation framework and used to investigate how cost of transport, stride length, stride frequency, and mechanical work performed by slow and fast twitch fibres were influenced by fibre type distribution across locomotion speeds (1.0 - 4.5 *m* · *s*^−1^). Cost of transport decreased as proportion of slow twitch fibres increased, while stride length and frequency were minimally affected by fibre type distribution during walking. During running, cost of transport decreased and increased with increasing proportions of slow twitch fibres above and below a threshold speed, respectively. Stride frequency decreased and stride length increased with increasing proportions of slow twitch fibres above this speed. These shifts in spatiotemporal characteristics may allow key muscles to operate close to their peak mechanical efficiency. In conclusion, fibre type distribution may influence both the energetics and biomechanics of gait in a locomotion speed dependent manner.

## 1 Introduction

Myosin molecules within muscle fibres convert chemical energy in nutrients to mechanical work through cross-bridge interactions with actin [1]. The biophysical properties of myosin isoforms influence the mechanical work that can be performed by each sarcomere, and this scales to whole muscle work that drives movements such as gait [[2]; [3]}. As myosin isoforms differ between fibre types, a muscle’s fibre type distribution affects both its contractile properties and the efficiency with which it converts chemical to mechanical energy [1,2,4]. As humans may self-optimize their gait to minimize the metabolic cost of travelling a given distance (cost of transport [COT]) [5,6], fibre type distribution is likely to influence whole-body COT both by altering optimal biomechanics for minimizing COT and by affecting muscle mechanical efficiency [2,3,7]. However, little is known about how muscle fibre type distribution influences whole-body biomechanics.

Muscle fibres are broadly classified as slow- or fast twitch depending on their twitch times. These fibre types predominantly express type I (slow) or type II (fast) myosin heavy chain isoforms, influencing their rate of adenosine triphosphate (ATP) hydrolysis [2,4]. Specifically, slow twitch muscle fibres hydrolyse ATP at a slower rate during slow contractions than fast twitch fibres [2,4]. This results in slower cross-bridge cycling, and consequently, slower maximal shortening velocities, lower force, and lower power output at a given submaximal shortening velocity in slow compared to fast twitch fibres [1,2,4]. Moreover, slow twitch fibres have higher mechanical efficiency at slow contraction speeds [1,2]. However, the rate of ATP hydrolysis increases more rapidly with increasing contraction velocities in slow compared to fast twitch fibres [2–4,8]. Consequently, fast twitch fibres may be more efficient at fast shortening velocities [1,2]. Human skeletal muscles contain both fibre types, but fibre type distributions vary substantially between muscles and individuals, and can change in response to use or disuse [3,7–10]. These differences have been proposed to influence whole-body movement, such as during gait [3,7,11], yet the evidence supporting this relationship remains unclear.

Most studies investigating the influence of fibre type distribution on locomotion have compared gait mechanics between groups with different fibre type profiles [7,11,12]. For example, Korhonen et al. [11] found that older runners had a lower proportion of fast fibres in the vastus lateralis, accompanied by lower peak push-off forces, shorter stride lengths, lower maximal speeds, longer contact times, and reduced knee extensor force-generating capacity compared with younger runners. Therefore, Korhonen et al. [11] suggested that the age-related decrease in fast twitch area fraction contributed to altered running biomechanics and muscular performance. Similarly, females generally have a higher proportion of slow twitch muscle fibres than males and also have lower stride lengths and higher stride frequencies [12]. However, as large morphological and physiological differences exist between these groups [11,12], the specific contribution of fibre type distribution to differences in running biomechanics cannot be determined. More recently, Swinnen et al. [7] compared physically active males with different triceps surae fibre type distributions and reported smaller speed-related increases in stride frequency and medial gastrocnemius activation, as well as smaller reductions in average net knee joint power, in individuals with more slow-twitch fibres. While their design better controlled for confounding factors, differences in cardiovascular fitness and training history may still have influenced their findings [7]. Moreover, fibre type distributions were quantified in only a small subset of locomotor muscles in all the aforementioned studies [7,11,12], limiting how well these results generalize to the entire locomotor system. Consequently, the isolated effects of fibre type distribution on gait biomechanics remains unclear, and *in vivo* investigation of these effects remains exceedingly difficult.

An attractive alternative is to use *in silico* musculoskeletal simulations, which allow individual muscle parameters to be manipulated in isolation. These simulations typically actuate rigid-link skeletal models using phenomenological Hill-type muscle models that capture the force-length and force-velocity properties of muscle [13–15]. Although computationally efficient, traditional Hill-type models represent an entire muscle with a single set of contractile parameters and a single time-varying activation state [15,16]. Consequently, they cannot represent the heterogeneity in contractile properties within or between muscles arising from differences in fibre type distribution, nor differential activation of slow- and fast twitch fibres [15,17]. A notable extension to the traditional Hill-type model uses two contractile elements with independently tuned parameters and independent activation states, enabling slow and fast fibre populations to be represented separately [18–21]. However, this muscle model has only been used in inverse simulations studies, investigating muscle fibre activation in a limited number of muscles [19,20]. As inverse simulations require experimentally measured kinematics and external forces as inputs, they cannot be used to investigate how changes in muscle fibre type distribution influence whole-body movement mechanics.

In contrast to inverse simulations, predictive simulations generate *de novo* movements from a musculoskeletal model and a performance criterion [22–24]. They are therefore well suited to investigate the influence of individual parameters, such as fibre type distribution, on whole-body biomechanics. For example, high-fidelity predictive simulations have been used to investigate the influence of musculoskeletal properties, such as plantar flexor strength and foot compliance, on gait biomechanics [23,24]. However, no study has used muscle models that represent the heterogeneity in muscle properties due to fibre type distribution in predictive simulations. The aim of the present study was to implement a muscle model that independently represents slow and fast muscle fibres in a predictive musculoskeletal simulation framework to investigate how variations in fibre type distribution influence gait biomechanics. Due to the greater efficiency of slow twitch fibres at slow contraction velocities and potential greater efficiency of fast fibres at high contraction velocities [2,4,7], we hypothesized that a larger proportion of slow fibres would reduce COT during at slow, but not faster, locomotion. Further, we hypothesized that a greater proportion of slow fibres would yield shorter stride lengths and higher stride frequency across all locomotion speeds. As a secondary aim, we explored the metabolic energy consumption, mechanical work performed, and mechanical efficiency of the slow and fast twitch fibres of the models across locomotion speeds.

## 2 Methods

### 2.1 Musculoskeletal model

We used a musculoskeletal model developed by D’Hondt et al. [24] based on previous OpenSim models [14,23] (Figure 1A). The model had 31 degrees of freedom (DOF; pelvis: 6 DOF, sacro-lumbar joint: 3 DOF, and hip: 3 DOF; knee: 1 DOF, ankle: 1 DOF, subtalar joint: 1 DOF, metatarsophalangeal joint: 1 DOF, glenohumeral joint: 3 DOF, and elbow: 1 DOF per side). The lower extremity and lumbo-sacral joints were actuated by 92 muscles, while the arms were actuated by eight idealized torque motors [23,24]. Non-linear rotational springs were used to represent the ligaments and capsules of each joint. Additionally, a small amount of damping (coefficient = 0.1 N · m · rad^−1^) was added at each joint. Foot-ground contact was modelled by five deformable spheres per foot interacting with a rigid plane, and the contact forces were calculated using a Hertzian stiffness and Hunt-Crossley dissipation model. The generic model was scaled to match the anthropometrics of one healthy female (mass = 62 kg, height = 1.70 m, age = 35 yrs.) based on data from Falisse et al. [23] using OpenSim’s scale tool [14].

**Figure 1:**
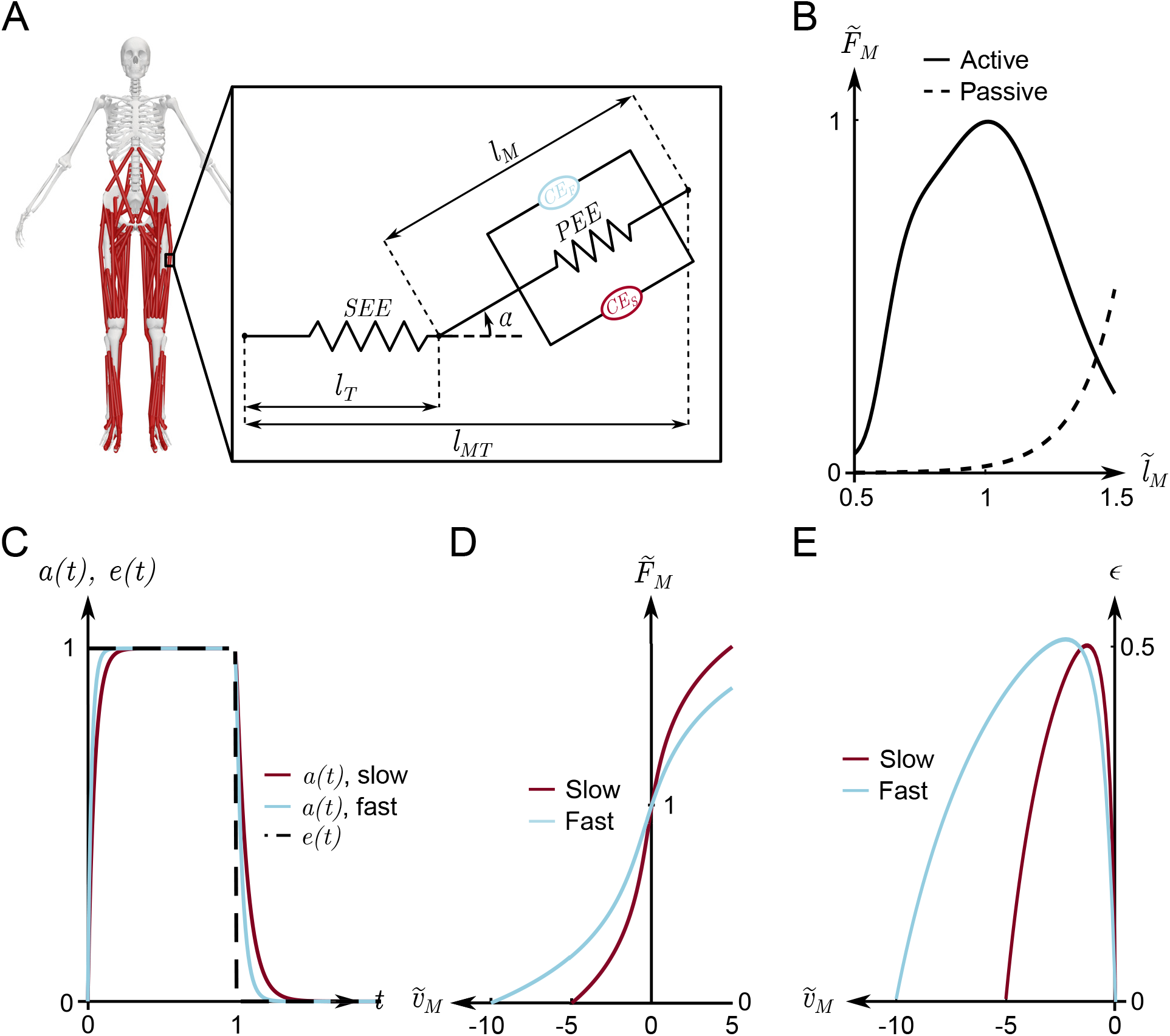
Illustration of the musculoskeletal model and a schematic of the muscle model used (panel A), as well as the force-length (panel B), excitation-activation (panel C), force-velocity (panel D), and efficiency-velocity (panel E) relationships of the muscle model. *SEE*: series elastic element, *PEE*: parallel elastic element, *CE*_*S/F*_ : slow and fast contractile elements, respectively, *l*_*M*_ : muscle length, *l*_*T*_ : tendon length, *l*_*MT*_ : muscle tendon unit length, *α*: pennation angle, 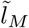: normalized muscle length, 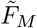: normalized muscle force, *a*: muscle activation, *e*: excitation, *t*: time, 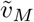: normalized muscle velocity, *ϵ*: mechanical efficiency

To independently represent the slow and fast twitch fibres of each muscle, the original Hill-type models were replaced with a two-element muscle model [20]. Briefly, the muscle was idealized as two length and velocity dependent parallel contractile elements, each representing one fibre type, arranged in parallel with a non-linear spring element (Figure 1). As the aggregate of contractile and parallel spring elements collectively represent all muscle fibres in the muscle belly, we refer to this as the muscle, rather than muscle fibre. The total force of the muscle (*F*_*M*_) was given by:

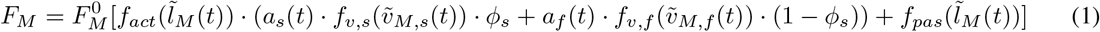

Where *t* is time, 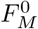 is maximal isometric force, *f*_*act*_ is a function describing the normalized active force developed at a given normalized muscle length 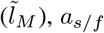 is the activation of slow and fast muscle fibres, *f*_*v,s/f*_ are functions expressing the normalized force developed by the slow and fast muscle fibres with respect to their normalized shortening velocity 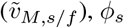 is the area fraction of slow twitch fibres, and *f*_*pas*_ expresses the normalized passive force as a function of normalized muscle length. The subscripts *s* and *f* are used to indicate the contractile element representing the slow and fast muscle fibres, respectively. While the normalized shortening velocities differed between the slow and fast contractile elements due to different maximal shortening velocities 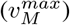, the absolute shortening velocity (*v*_*M*_ (*t*)) of both contractile elements was the same. The contractile and parallel elastic elements were arranged in series with a non-linear spring representing the tendon at an angle *α*, representing the pennation angle. For each muscle tendon unit (MTU), the relationships between the tendon and muscle are governed by the following equations [15]:

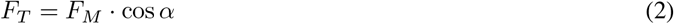

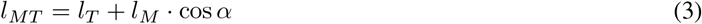

Where *F*_*T*_ is tendon force, *l*_*MT*_ is MTU length, *l*_*T*_ is tendon length, and *l*_*M*_ is muscle length (see Figure 1 for muscle model geometry). The length and velocity dependence of the force produced by the contractile and elastic elements were modelled as described by De Groote et al. [15] (Figure 1). To represent the different properties of slow and fast twitch fibres, maximal shortening velocities of 5 and 10 optimal muscle fibre lengths per second were used for the slow and fast contractile elements, respectively [18,20]. Additionally, each fibre’s excitation-activation relationship was modelled with a transfer function [17,25], and activation time constants (*τ*_*act*_) of 45 ms and 25 ms were used for slow and fast contractile elements, respectively [20]. The deactivation time constants (*τ*_*deact*_) were given by *τ*_*deact*_ = *τ*_*act*_ · 0.6^−1^ [16,20]. Lastly, the area fraction of slow and fast twitch fibres in the muscles of the nominal model were taken from Uchida et al. [26]. Muscles for which no experimental fibre type data were available were assumed to have an equal proportion of slow and fast muscle fibres [26].

### 2.2 Predictive simlation framework

Predictive simulations of walking and running were formulated as optimal control problems (OCP) [22,23]. The OCPs were solved to find the control (**u**) and corresponding state (**x**) trajectories that minimized the cost functional (*J*), subject to dynamic constraints enforcing Newtonian system dynamics (i.e.,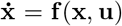, task constrains enforcing left-right symmetry and average forward gait speed, and path constraints preventing limb collision [22,23]. We used a multi-objective cost functional that minimized the time integral of a weighted sum of metabolic rate (**Ė**), muscle activations (**a**), the second time derivative of generalized joint coordinates 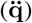, passive joint torques (*τ*_*pass*_), and excitation of idealized torque actuators driving the arms (**e**_*arms*_).

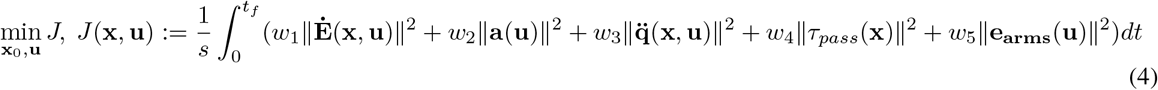

where **x**_0_ is the initial state, *s* is the distance travelled, *t*_*f*_ is the final time, *w*_1−5_ are weight factors, and ∥.∥ indicates the Euclidean norm. Metabolic energy rate was calculated with a smooth, continuous metabolic function [24,27,28], which was modified to use the activations of the muscles’ slow and fast fibres, rather than the orderly recruitment imposed in the original model [27,28].

The PredSim framework for predictive musculoskeletal simulations was used to solve the OCP [22–24]. Briefly, the OCP was formulated in MATLAB (R2024b, MathWorks, Natick, MA, USA) and used a direct collocation scheme with a cubic Radau quadrature over 100 mesh intervals to transcribe the OCP to a non-linear program (NLP) [23,24]. The system dynamics were formulated implicitly and derivatives required for gradient based optimization were calculated using algorithmic differentiation in CasADi 3.7.1 [22,29]. The NLP problem was solved using the IPOPT and MUMPS solvers [30,31]. Each simulation was performed using both a quasi-random initial guess and the nearest converged solution as initial guess, retaining only the solution with the lowest cost [23]. For the speeds closest to the walk-run transition (2.0 m/s and 2.5 m/s, see below for details), a quasi-random initial guess, as well as the nearest converged walking and running solutions were used as initial guesses [23]. All simulations were run on a laptop (MSI Stealth AI16, Intel Core Ultra 9 185H CPU, 32GB RAM), allocating two CPU threads to each simulation run in parallel.

### 2.3 Model validation

The predicted joint angle, joint torque, joint power, muscle activation, and ground reaction force time-series from simulated gait at 1.33 m · s^−1^, 3.0 m · s^−1^, and 4.0 m · s^−1^ using the nominal fibre type distribution were compared to experimental data from Falisse et al. [23] (1.33 m · s^−1^) and Hamner and Delp [13] (3.0 m · s^−1^ and 4.0 m · s^−1^) for validation. Specifically, predicted ankle dorsiflexion, knee extension, and hip flexion, adduction, and internal rotation angles, torques, and powers, as well as gluteus medius, semimembranosus, vastus lateralis, rectus femoris, tibials anterior, medial gastrocnemius, lateral gastrocnemius, and soleus activations, and ground reaction forces were qualitatively compared to corresponding experimental data. These specific variables were selected for comparison as they were available in both experimental data sets [13,23]. As each muscle had separate activation for the slow and fast contractile elements, their sum weighted by muscle fibre area fractions were compared to the experimental eletromyography (EMG) amplitudes at each time step. Additionally, normalized cross-correlation coefficients (R) were calculated between simulated and mean experimental joint angles, joint torques, and joint powers to quantify the similarities in signal shape [24,32]. The similarity between time-series were considered strong if R was greater than 0.7, moderate if R was between 0.5 and 0.7, fair if R was between 0.3 and 0.5, and poor if R was less than 0.3 [32]. We also computed the standard deviations (SD)-weighted root mean square errors (RMSE) for joint angles, torques, and powers [24].

### 2.4 Influence of fibre type distribution

To investigate the influence of fibre type distribution on simulated gait mechanics, we varied the model’s fibre-type distribution from 6% to 96% slow fibres on average across all muscles (see Figure 2 for muscle-specific fibre-type distributions). Simulations were generated for 11 gait speeds between 1.0 and 4.5 m · s^−1^, and classified as walking or running based on whether centre of mass kinetic and potential energies varied in anti-phase or phase, respectively [23]. Predicted COT, stride length, and stride frequency were extracted for comparison. COT was calculated by integrating muscle metabolic rates over the gait cycle, summing across muscles, adding a basal metabolic rate, and dividing by body mass and distance travelled [23,27]. Stride length was taken to be the forward distance travelled by the pelvis centre of mass over one stride, while stride frequency was computed as the inverse of the stride duration. Relationships between speed and COT were characterized by fitting linear and quadratic polynomials to the data, and the fits were evaluated using coefficients of determination and normalized residuals. Quadratic models provided the best fit for all variables during walking and running (online supplement tables S1-S6). The walking and running speeds that minimized COT for each fibre type distribution were estimated from the minima of the fitted quadratic functions. To examine fibre type specific energetics, metabolic energy consumption, positive mechanical work, and mechanical efficiency were calculated separately for slow and fast contractile elements. Specifically, metabolic energy consumption was obtained by integrating the contractile elements’ metabolic rates over the gait cycle and summing the integrals across all muscles. As the metabolic model only accounted for positive work performed by the muscle, the mechanical work performed by slow and fast contractile elements was calculated by integrating the positive work performed over the gait cycle and summed across muscles. Mechanical efficiency was defined as the ratio of mechanical work to metabolic energy consumption. These variables were evaluated across all locomotion speeds for three representative fibre type distributions (6% slow, nominal, and 96% slow). Because the simulations produced deterministic theoretical predictions, null-hypothesis statistical testing was not performed.

**Figure 2:**
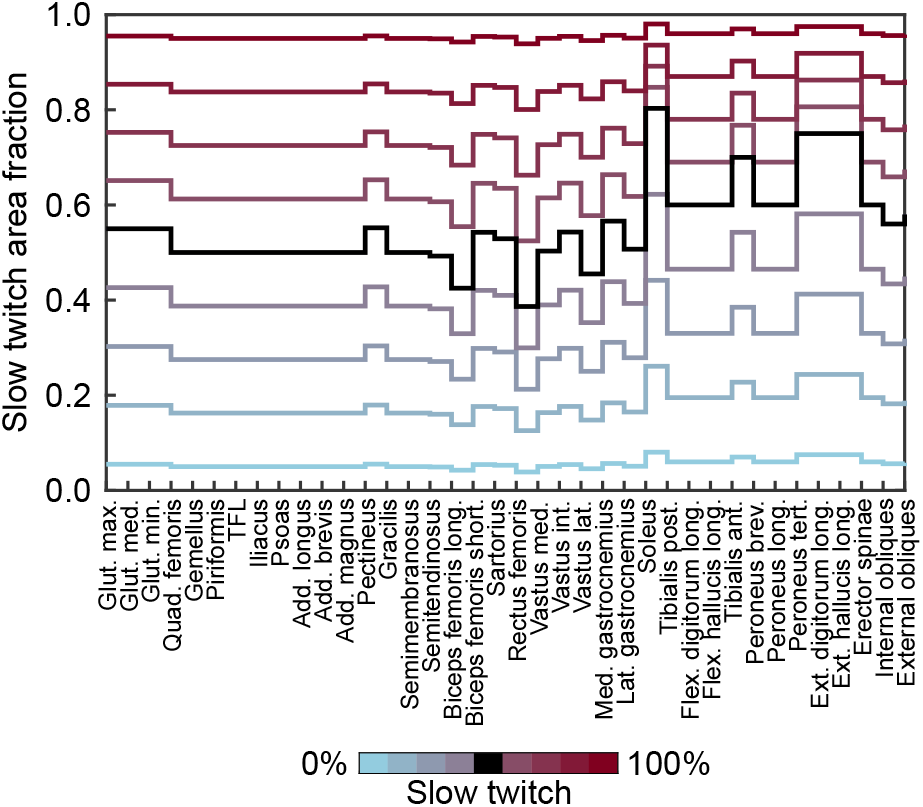
The fibre type distributions of the nine models that were used in the present study. The black line indicates the fibre type distribution of the nominal model. Each line represents the model whose average slow twitch area fraction corresponds to the colour depicted in the scale.

## 3 Results

### 3.1 Model validation

All simulations at speeds below 2.5 m · s^−1^ resulted in walking, while the remaining simulations resulted in running. Most hip, knee, and ankle angles, torques, and powers from the simulations at 1.33 m · s^−1^ were qualitatively similar to experimental time-series (online supplement figures S1). The simulated time-series had strong cross-correlations with the experimental means for 10 out of 15 joint coordinates, torques, and powers (R = 0.85 to R = 0.99; online supplement table S7). However, peak ankle dorsiflexion and plantar flexion were visibly smaller in simulated compared to experimental results (online supplement figures S1). The cross-correlations were moderate for the ankle angle (*R* = 0.66), knee power (R = 0.69), and hip adduction power (R = 0.67). Further, hip internal/external rotation angle and power time-series had different shape than experimental time-series, and both hip internal/external rotation angle (R = 0.16) and power (R = 0.23) exhibited poor cross-correlation with experimental data. RMSEs ranged from 1.63 to 7.34 SDs (see discussion for likely explanation; online supplement table S8).

All joint angles, torques, and powers for simulations at 3.0 m · s^−1^ were qualitatively similar to the experimental data, except for smaller peak knee flexion during the stance and swing phases of simulated running (online supplement figures S2). All joint angles, torques, and powers except hip internal/external rotation angle had strong cross-correlations with experimental data (R = 0.74 to R = 0.99; online supplement table S9). The hip internal/external rotation angles had moderate (R = 0.59) cross-correlation with experimental data. RMSEs ranged from 0.80 to 3.27 SDs (online supplement table S10).

Hip, knee, and ankle angles, torques, and powers during running simulations at 4.0 m · s^−1^ were also qualitatively similar to the experimental time-series (online supplement figures S3). Further, all joint angles, torques, and powers except hip internal/external rotation angle, torque, and power had strong cross-correlations with the experimental data for simulation at 4.0 m · s^−1^ (R = 0.79 to 0.99; online supplement table S11). However, hip internal/external rotation angle (R = 0.59) and torque (R = 0.63) had moderate, while hip internal/external rotation power (R = 0.35) had poor cross-correlations with the experimental trials, respectively. RMSEs ranged from 0.85 to 3.92 SDs (online supplement table S12). The simulations captured the increase in hip flexion/extension range of movement, peak hip flexion and extension moments, peak knee extension moment, peak ankle plantar flexion moment, as well as increases in peak breaking, propulsive, and vertical ground reaction forces observed with increasing speed in the experimental running data [13]. However, the increases in peak knee flexion during swing and peak ankle plantar flexion during stance were not captured by the simulations.

Qualitatively, deactivation of the gluteus medius and tibialis anterior muscles was slightly delayed in simulated walking, while rectus femoris activations did not reproduce the experimentally observed EMG well for either gait (online supplement figures S1, S2, S3). Remaining muscle activation, as well as ground reaction force time-series were qualitatively similar to experimental time-series.

### 3.2 Walking

COT had a quadratic relationship with walking speed for all fibre type distributions (Figure 3). COT in the model with the largest proportion of slow fibres was 78.6 *±* 1.8% of the COT in the model with the smallest proportion of slow twitch fibres on average (*±*SD) across walking speeds. Models with intermediate area fractions of slow twitch fibres had intermediate COT for all walking speeds. Stride length increased from 1.267 *±* 0.003 m to 1.536 *±* 0.015 m as speed increased from 1.0 m · s^−1^ to 1.5 m · s^−1^ and subsequently plateaued, with stride lengths of 1.543 *±* 0.014 m and 1.537*±*0.011 m at 1.75 and 2.0 m·s^−1^, respectively. Stride frequencies increased nearly linearly from 0.789*±*0.002 Hz to 1.301 *±* 0.009 Hz as speed increased from 1.0 m · s^−1^ to 2.0 m · s^−1^ (Figure 3). However, stride lengths and stride frequencies differed minimally between different fibre type distributions with the largest differences being 0.056 m for stride length and 0.035 Hz for stride frequency, both at 1.5 m · s^−1^.

**Figure 3:**
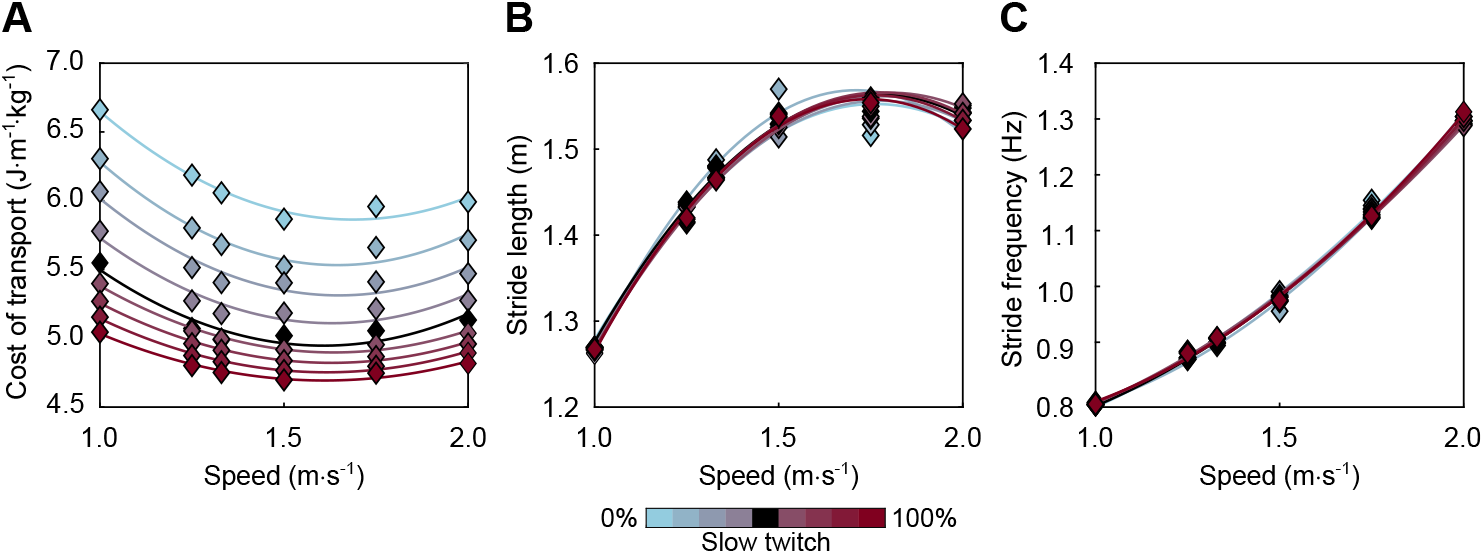
The cost of transport (COT; panel A), stride length (panel B), and stride frequency (panel C) during walking at speeds from 1.0 m · s^−1^ to 2.0 m · s^−1^ with different fibre type distributions. The nominal model is illustrated in black, while red and blue illustrate models with various levels of increased or decreased proportions of slow twitch fibres, respectively.

The walking speeds that were estimated to minimize COT were slowest (1.61 m · s^−1^) in the model with the largest proportion of slow twitch fibres and fastest (1.69 m · s^−1^) in the model with the smallest proportion of slow twitch fibres (Figure 4).

**Figure 4:**
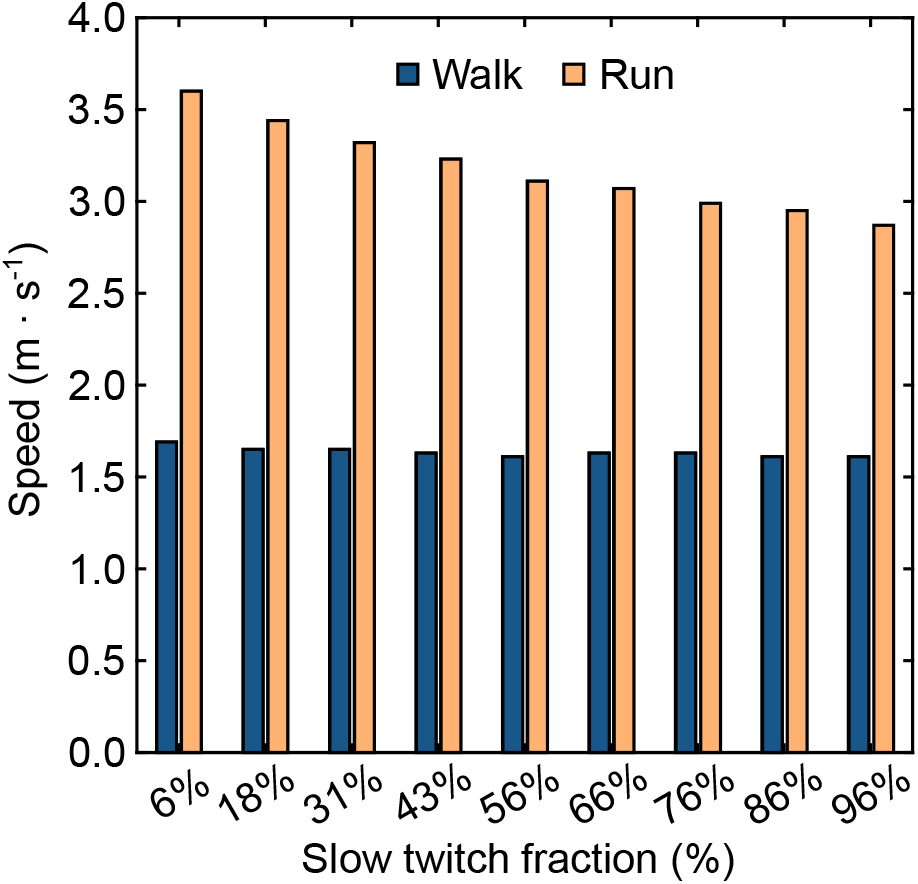
Walking and running speeds that are estimated to minimize the metabolic cost of transport (COT) for each average slow twitch area fraction. The estimates are based on finding the minimum of quadratic functions fit to the COT and speed data.

Absolute metabolic energy consumption increased with walking speed in the 6% slow twitch, nominal, and 96% slow twitch conditions (Figure 5). The mechanical work performed followed a similar trend to that of the metabolic energy consumption. However, the mechanical work increased at a higher rate than the increase in metabolic energy in all but the slow twitch fibres in the 6% slow twitch condition. Consequently, mechanical efficiency increased with increasing walking speed in all except the slow twitch fibres in the 6% slow twitch condition.

**Figure 5:**
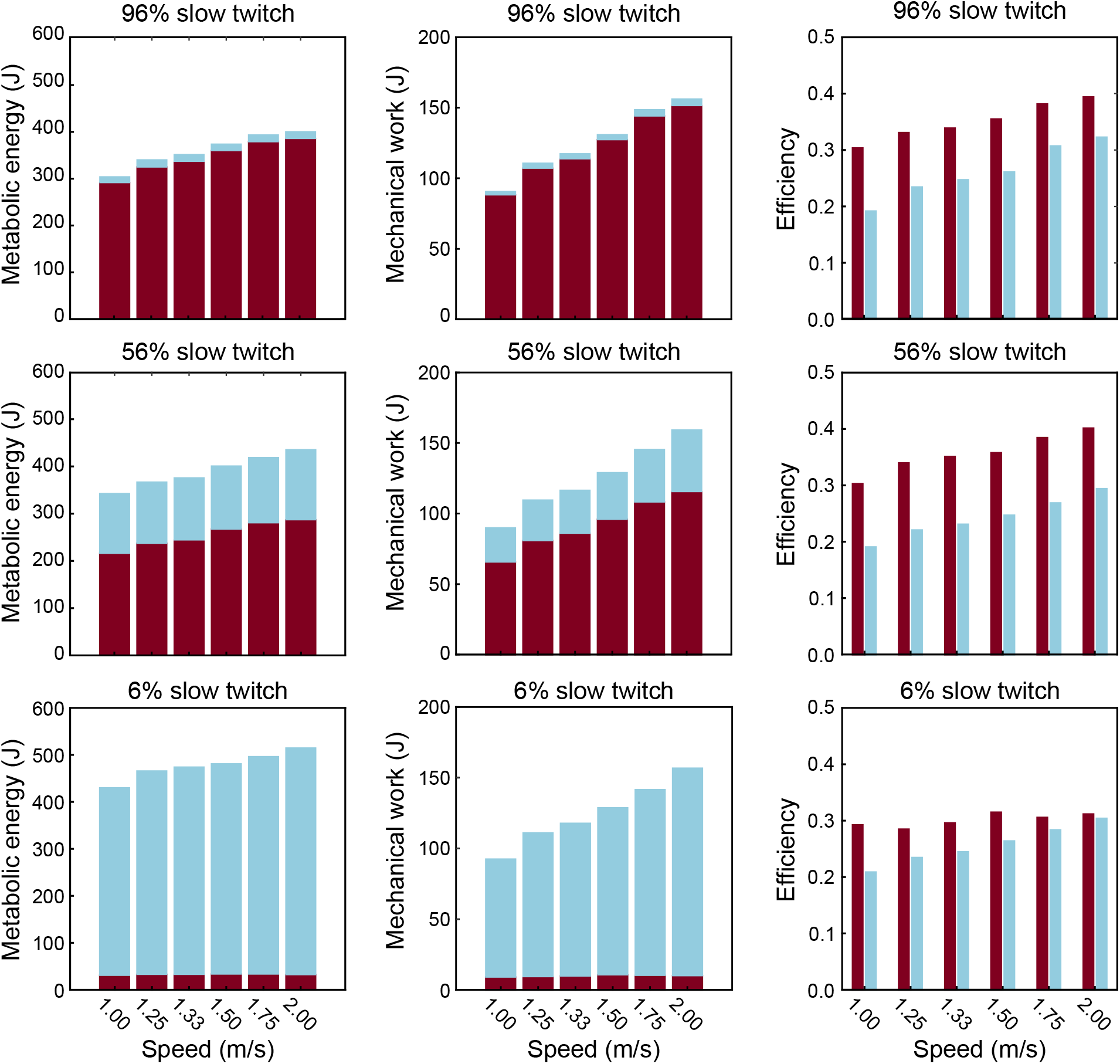
Metabolic energy consumption, mechanical work, and mechanical efficiency over one stride for all slow (light blue) and fast (red) twitch fibres across walking speeds in the 6% slow twitch (bottom panels), 56% slow twitch (nominal; middle panels), and 96% slow twitch (top panels).

### 3.3 Running

As in walking, COT had a quadratic relation to running speed for all fibre type distributions (Figure 6). However, there was an interaction effect between fibre type distribution and COT in running. This resulted in COT being lowest in the model with 96% (4.00 J · kg^−1^ · min^−1^, 3.95 J · kg^−1^ · min^−1^, and 4.08 J · kg^−1^ · min^−1^, respectively) and highest in the model with 6% (4.99 J · kg^−1^ · min^−1^, 4.70 J · kg^−1^ · min^−1^, and 4.61 J · kg^−1^ · min^−1^, respectively) slow twitch fibres at speeds of 2.5, 3.0, and 3.5 m · s^−1^. The nominal model had the lowest COT (4.41 J · kg^−1^ · min^−1^) at 4.0 m · s^−1^ and COT was highest in the model with 6% slow twitch fibres (4.66 J · kg^−1^ · min^−1^) at this speed. However, at 4.5 m · s^−1^ COT was lowest in the model with 43% slow twitch fibres (4.74 J · kg^−1^ · min^−1^) and highest in the model with 86% slow twitch fibres (4.97 J · kg^−1^ · min^−1^). Stride length increased nearly linearly from 1.431 *±* 0.003 m to 1.991 *±* 0.013 m with increases in speed from 2.5 m · s^−1^ to 4.0 m · s^−1^. However, at 4.5 m · s^−1^ the stride length of models with different fibre type distributions diverged considerably (Figure 6). Specifically, the model with the largest proportion of slow twitch fibres took strides that were 0.154 m longer that those taken by the model with the smallest proportion of slow twitch fibres. The remaining models varied between the 6% and 96% slow twitch fibre models in the order of their slow twitch fibre proportions (Figure 6). In contrast to stride lengths, stride frequencies increased from 1.75 *±* 0.00 Hz at 2.5 m · s^−1^ to 2.01 *±* 0.01 Hz at 4.0 m · s^−1^, and plateaued thereafter. At the 4.5 m · s^−1^ speed, stride frequencies were higher in the model with 6% slow twitch fibres (2.08 Hz) and lower in the model with 96% slow twitch fibres (1.94 Hz). Stride frequencies of the remaining models were distributed between these in order of their slow twitch area fractions (Figure 6).

**Figure 6:**
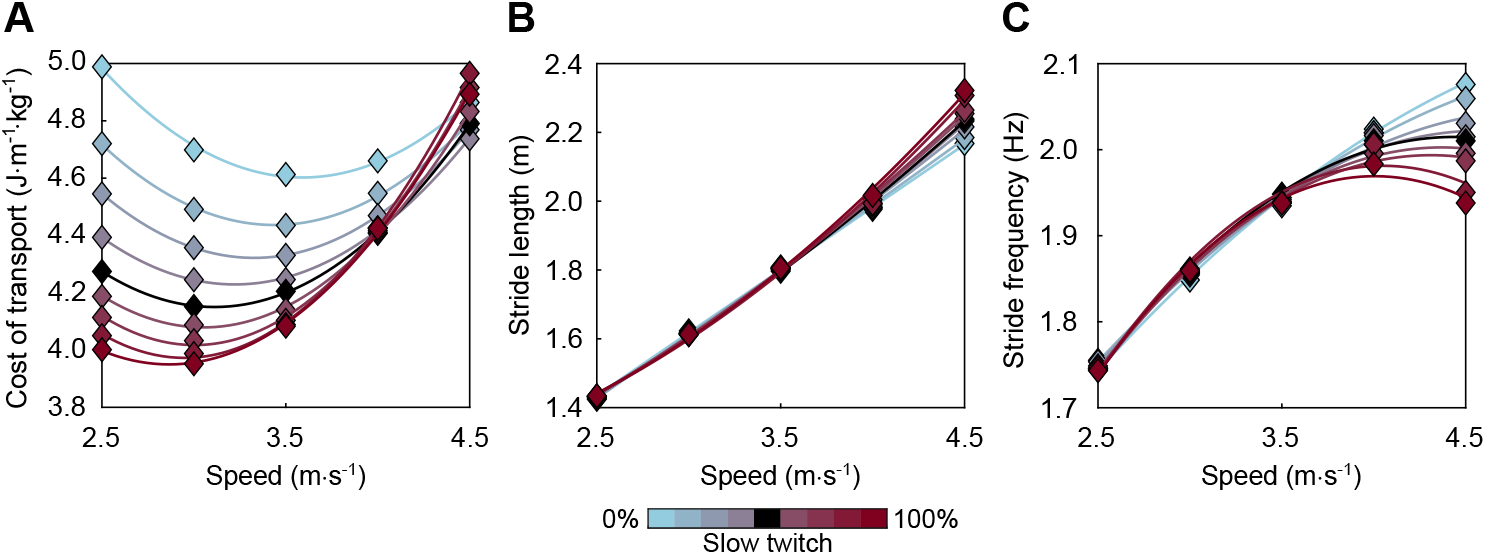
The cost of transport (COT; panel A), stride length (panel B), and stride frequency (panel C) during running at speeds from 2.5 m · s^−1^ to 4.5 m · s^−1^ with different fibre type distributions. The nominal model is illustrated in black, while red and blue illustrate models with various levels of increased or decreased proportions of slow twitch fibres, respectively.

The running speeds estimated to minimize COT were higher in models with a larger proportion of fast twitch fibres and lower in models with a larger proportion of slow twitch fibres (Figure 4).

Lastly, absolute metabolic energy consumption increased with running speed in the 6% slow twitch, nominal, and 96% slow twitch conditions (Figure 7). The rate of increase with respect to speed was greater in fast compared to slow twitch fibres. Similar to metabolic energy consumption, the mechanical work increased from slower to faster running speeds, with greater rates of increase with respect to speed in fast compared to slow fibres. The mechanical efficiency increased with increasing running speed in both fibre types across all fibre type distributions. However, mechanical efficiency increased more in fast compared to slow fibres from running speeds of 2.5 m · s^−1^ to 4.5 m · s^−1^.

**Figure 7:**
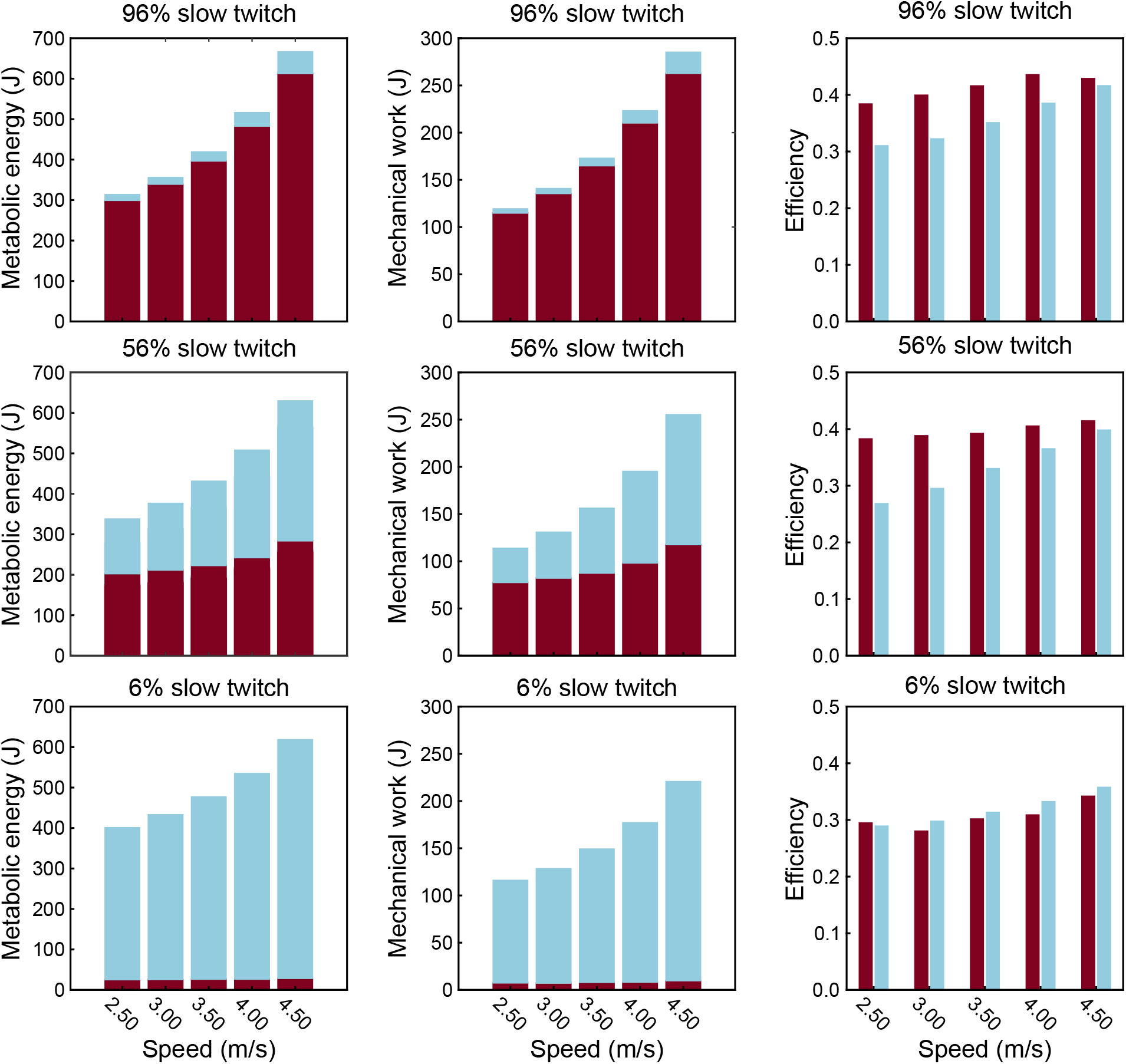
Metabolic energy consumption, mechanical work, and mechanical efficiency over one stride for all slow (light blue) and fast (red) twitch fibres across running speeds in the 6% slow twitch (bottom panels), 56% slow twitch (nominal; middle panels), and 96% slow twitch (top panels).

## 4 Discussion

In the present study, we implemented a muscle model that independently represents slow and fast twitch muscle fibres within a predictive simulation framework. This enabled us to investigate how fibre type distribution, through its effects on muscle fibre properties, scales to influence whole-body gait energetics and mechanics. Overall, the simulations resulted in plausible walking and running biomechanics compared to experimental studies [13,23]. In support of our hypotheses, COT decreased as the proportion of slow twitch fibres increased during walking. In running, the proportion of slow twitch fibres interacted with running speed, with models having larger proportions of fast-twitch fibres exhibiting lower COT at the fastest running speed. Fibre type distribution also influenced spatiotemporal gait characteristics at the fastest running speeds, while stride lengths and frequencies were largely unaffected at speeds below 4.0 m · s^−1^. Contrary to our hypotheses, models with greater proportions of fast twitch fibres had higher stride frequencies at the fastest running speed. Although these results provide novel insight into how fibre type distribution may influence gait mechanics, they should be interpreted as theoretical predictions requiring experimental corroboration.

The model with the nominal fibre type distribution produced physiologically plausible gait across all speeds, and generally compared well with experimental walking and running data [13,23]. Most simulated joint coordinates, torques, and powers exhibited strong cross-correlations (R > 0.7) with experimental data. Hip internal/external rotation angle and power were notable exceptions, showing poor cross-correlations with experimental data during both walking and running. Further, RMSEs were frequently large (RMSE > 3 SD) when compared to the experimental data from walking. While these RMSEs exceeded those previously reported [32], the data set from Falisse et al. [23] included only one individual. Thus, this data set contained SDs reflecting within-individual variability, which is considerably smaller than between-individual variability [33]. Consequently, large RMSEs were not surprising for walking [23]. In contrast, RMSEs for running, which were weighted by the larger between-individual SDs [13], were substantially smaller. Together these observations suggest that large RMSEs in walking largely reflects the use within-individual SDs for normalization, rather than poor simulation performance. While the predictive simulations did not capture all aspects of experimental gait, the simulations do represent plausible walking and running.

The predicted COT displayed a quadratic relationship with gait speed in both walking and running gaits, as observed in previous studies [34–37]. COT decreased with larger proportions of slow twitch fibres during walking, while an interaction between the proportion of slow twitch fibres and gait speed was predicted during running. Specifically, models with larger proportions of slow twitch fibres had lower COT at running speeds of 2.5 to 4.0 m · s^−1^, but higher COT at 4.5 m · s^−1^. While previous experimental results on the relationship between fibre type distribution and COT are conflicting, several studies report lower COT in individuals with a higher proportion of slow twitch fibres when running at slow speeds, and higher COT at faster speeds [3,7,38–41]. For example, Swinnen et al. [7] found that individuals with a larger proportion of slow twitch fibres in the gastrocnemius and soleus had lower COT during running at speeds between 2.0 and 4.0 m · s^−1^. In contrast, Pellegrino et al. [40] found that a larger proportion of slow twitch fibres in the vastus lateralis was associated with higher COT when running at 4.56 m · s^−1^. Taken together, previous experimental studies and our simulations suggest that a threshold running speed exists above which a larger proportion of fast twitch fibres becomes metabolically advantageous.

This interpretation is consistent with the metabolic and contractile properties of individual muscle fibres. Specifically, fast twitch fibres can perform work at higher rates with greater efficiency than slow fibres at high shortening velocities [1,2]. Therefore, it has been proposed that fast fibres may be recruited prior to slow fibres during fast muscle contractions to minimize metabolic cost [19,21]. While some suggest that major locomotion muscles, such as vastus lateralis, and medial and lateral gastrocnemius, operate at slow speeds [42–45], the shortening velocity of other important locomotor muscles increases considerably with increasing gait speed [45,46]. Our simulations predicted increasing active shortening velocities of multiple large locomotor muscles with increasing gait speed. Some muscles, such as the vasti, lateral gastrocnemius, and soleus muscles, actively shortened at speeds where fast twitch fibres are more efficient than slow twitch (online supplement figure S4) [1,2]. This behaviour was also reflected in the predicted efficiency of the slow and fast twitch fibres, which increased at similar rates in walking but increased more rapidly in fast twitch fibres during running. Thus, models with a larger pool of fast twitch fibres to recruit from may have gained a metabolic advantage at the fastest running speeds.

Contrary to our hypotheses, stride lengths increased while stride frequency decreased as the proportion of slow twitch fibres increased during running. These effects were negligible during walking and slower running, but became substantial at the fastest running speeds. One potential explanation is that the muscles involved in propelling the body were able to perform the required work in a shorter amount of time in the models with a larger proportion of fast twitch fibres. In contrast, we had hypothesized that less work would be performed during stance with a larger proportion of slow twitch fibres, thereby reducing stride length and increasing stride frequency. Nonetheless, the increase in stride frequency with more fast twitch fibres agrees with the findings of Swinnen et al. [7], who observed that stride frequency increased at a higher rate in individuals with larger proportions of fast twitch fibres in the gastrocnemius and soleus muscles. Interestingly, fatigue during constant-speed running has also been associated with longer stride lengths and lower stride frequencies [47]. Because fast twitch fibres are more fatigable than slow twitch fibres, fast fibres would likely fatigue first, resulting in greater reliance on slow twitch fibres. The present findings therefore offer a potential mechanism underlying fatigue-related changes in running mechanics.

Despite providing novel insights into the influence of fibre type distribution on gait mechanics, this study is not without limitations. First, COT during walking was overestimated compared experimental observations [48], whereas COT in running were comparable experimental results [37]. A potential explanation is that metabolic models derived from individual muscle fibres do not fully capture how metabolic energy expenditure scale to whole muscles. Alternatively, inaccuracies in the musculoskeletal or ground contact models may have contributed to the discrepancy between simulated and experimental walking [49]. Nonetheless, the general relationship between COT and speed, and its response to fibre type distribution were consistent with the literature [3,7,34,35]. Thus, the metabolic model appears to have adequately represented muscle energetics for addressing the aims of this study. Nonetheless, the COT magnitudes and the predicted relationship between gait speed and COT should be interpreted with caution. Another limitation is that stride frequencies during running were higher than those typically observed in experiments [7,11]. As stride frequency tend to increase with running speed, some of the predicted effects of fibre type distribution on stride length and frequency may only manifest at higher running speeds. More broadly, all our findings should be treated as theoretical predictions, as the simulations were not driven by experimental data. As with all computer models, the results reflect a trade-off between model complexity and computational tractability, dependent on the underlying modelling assumptions and the chosen cost function [19]. Different modelling choices or objective functions may therefore yield different results.

In conclusion, our simulations predict that muscle fibre type distribution influences both the energetics and mechanics of gait. A greater proportion of slow twitch fibres may reduce metabolic cost at slow gait speeds, whereas a threshold speed appears to exist above which a larger proportion of fast twitch fibres lowers COT. This shift may arise because faster running favours higher stride frequencies, and consequently, muscle shortening velocities, allowing fast twitch fibres to operate closer to their peak efficiency.

## 5 Acknowledgements

The authors express their gratitude to Dr. Nilima Nigam for insightful discussions about handling numerically unstable cases and for feedback on the manuscript. We also acknowledge the support of the Government of Canada’s New Frontiers in Research Fund (NFRF), [NFRFE-2023-00166].

## 6 Data Accessibility

All simulation results are available from https://doi.org/10.5281/zenodo.22236564 while source code can be obtained from https://github.com/TorsteinDaehlin/predsim-multifibre/tree/multifibre-release.

