## Supplemental file for "The effects of muscle fibre type distribution on gait biomechanics: A predictive simulation study"

**Online supplement**

Torstein E. Dæhlin Stephanie A. Ross Friedl De Groote
James M. Wakeling

2025-11-17

### Introduction

This document contains supplementary material for a research study where we used predictive musculoskeletal simulations to investigate the effects of muscle fibre type distribution on cost of transport and gait biomechanics:

– REFERENCE TO PREPRINT/PUBLISHED ARTICLE HERE –

See manuscript for further details.

### Cost of transport, stride length, and stride frequency polynomial goodness of fit

The goodness of fit of linear and quadratic polynomials to the cost of transport, stride length, and stride frequency results during walking and running were assessed by the normalized residuals and coefficients of determination. These measures of goodness of fit are presented in tables Table S 1, Table S 2, Table S 3, Table S 4, Table S 5, and Table S 6.

#### Walking

Table S 1: Normalized residuals and coefficients of determination ($R^2$) for linear and quadratic polynomial fits to cost of transport during simulated walking at speeds from 1.0 to 2.0 m/s.

| Slow twitch fraction | Residuals - Linear | $R^2$ - Linear | Residuals - Quadratic | $R^2$ - Quadratic |
|---|---|---|---|---|
| 6% | 0.428 | 0.551 | 0.112 | 0.969 |
| 18% | 0.468 | 0.412 | 0.133 | 0.952 |
| 31% | 0.444 | 0.412 | 0.156 | 0.927 |
| 43% | 0.421 | 0.328 | 0.148 | 0.916 |
| 56% | 0.403 | 0.249 | 0.153 | 0.892 |
| 66% | 0.304 | 0.386 | 0.055 | 0.980 |
| 76% | 0.277 | 0.363 | 0.040 | 0.987 |
| 86% | 0.259 | 0.311 | 0.035 | 0.987 |
| 96% | 0.233 | 0.283 | 0.045 | 0.974 |

Table S 2: Normalized residuals and coefficients of determination ($R^2$) for linear and quadratic polynomial fits to stride length during simulated walking at speeds from 1.0 to 2.0 m/s.

| Slow twitch fraction | Residuals - Linear | $R^2$ - Linear | Residuals - Quadratic | $R^2$ - Quadratic |
|---|---|---|---|---|
| 6% | 0.128 | 0.685 | 0.044 | 0.963 |
| 18% | 0.163 | 0.597 | 0.047 | 0.964 |
| 31% | 0.122 | 0.734 | 0.031 | 0.982 |
| 43% | 0.128 | 0.699 | 0.023 | 0.990 |
| 56% | 0.123 | 0.731 | 0.026 | 0.988 |
| 66% | 0.117 | 0.776 | 0.023 | 0.991 |
| 76% | 0.126 | 0.741 | 0.020 | 0.993 |
| 86% | 0.129 | 0.719 | 0.018 | 0.995 |
| 96% | 0.133 | 0.692 | 0.016 | 0.996 |

Table S 3: Normalized residuals and coefficients of determination ($R^2$) for linear and quadratic polynomial fits to stride frequency during simulated walking at speeds from 1.0 to 2.0 m/s.

| Slow twitch fraction | Residuals - Linear | $R^2$ - Linear | Residuals - Quadratic | $R^2$ - Quadratic |
|---|---|---|---|---|
| 6% | 0.063 | 0.979 | 0.025 | 0.997 |
| 18% | 0.087 | 0.961 | 0.027 | 0.996 |

| Slow twitch fraction | Residuals - Linear | $R^2$ - Linear | Residuals - Quadratic | $R^2$ - Quadratic |
|---|---|---|---|---|
| 31% | 0.058 | 0.981 | 0.014 | 0.999 |
| 43% | 0.063 | 0.978 | 0.009 | 1.000 |
| 56% | 0.059 | 0.980 | 0.011 | 0.999 |
| 66% | 0.056 | 0.981 | 0.015 | 0.999 |
| 76% | 0.063 | 0.977 | 0.016 | 0.999 |
| 86% | 0.066 | 0.975 | 0.015 | 0.999 |
| 96% | 0.070 | 0.973 | 0.014 | 0.999 |

### Running

Table S 4: Normalized residuals and coefficients of determination ($R^2$) for linear and quadratic polynomial fits to cost of transport during simulated running at speeds from 2.5 to 4.5 m/s.

| Slow twitch fraction | Residuals - Linear | $R^2$ - Linear | Residuals - Quadratic | $R^2$ - Quadratic |
|---|---|---|---|---|
| 6% | 0.300 | 0.084 | 0.019 | 0.996 |
| 18% | 0.298 | 0.042 | 0.014 | 0.998 |
| 31% | 0.305 | 0.252 | 0.001 | 1.000 |
| 43% | 0.293 | 0.467 | 0.011 | 0.999 |
| 56% | 0.310 | 0.633 | 0.007 | 1.000 |
| 66% | 0.338 | 0.695 | 0.015 | 0.999 |
| 76% | 0.376 | 0.737 | 0.021 | 0.999 |
| 86% | 0.386 | 0.776 | 0.018 | 1.000 |
| 96% | 0.333 | 0.821 | 0.022 | 0.999 |

Table S 5: Normalized residuals and coefficients of determination ($R^2$) for linear and quadratic polynomial fits to stride length during simulated running at speeds from 2.5 to 4.5 m/s.

| Slow twitch fraction | Residuals - Linear | $R^2$ - Linear | Residuals - Quadratic | $R^2$ - Quadratic |
|---|---|---|---|---|
| 6% | 0.009 | 1.000 | 0.009 | 1.000 |
| 18% | 0.012 | 1.000 | 0.011 | 1.000 |
| 31% | 0.030 | 0.998 | 0.024 | 0.998 |
| 43% | 0.039 | 0.996 | 0.024 | 0.999 |
| 56% | 0.043 | 0.995 | 0.018 | 0.999 |

| Slow twitch fraction | Residuals - Linear | $R^2$ - Linear | Residuals - Quadratic | $R^2$ - Quadratic |
|---|---|---|---|---|
| 66% | 0.048 | 0.994 | 0.022 | 0.999 |
| 76% | 0.047 | 0.995 | 0.017 | 0.999 |
| 86% | 0.080 | 0.986 | 0.040 | 0.997 |
| 96% | 0.077 | 0.988 | 0.027 | 0.998 |

Table S 6: Normalized residuals and coefficients of determination ($R^2$) for linear and quadratic polynomial fits to stride frequency during simulated running at speeds from 2.5 to 4.5 m/s.

| Slow twitch fraction | Residuals - Linear | $R^2$ - Linear | Residuals - Quadratic | $R^2$ - Quadratic |
|---|---|---|---|---|
| 6% | 0.032 | 0.985 | 0.006 | 0.999 |
| 18% | 0.036 | 0.979 | 0.008 | 0.999 |
| 31% | 0.051 | 0.952 | 0.021 | 0.992 |
| 43% | 0.063 | 0.923 | 0.020 | 0.992 |
| 56% | 0.069 | 0.907 | 0.013 | 0.997 |
| 66% | 0.072 | 0.891 | 0.017 | 0.994 |
| 76% | 0.072 | 0.880 | 0.012 | 0.997 |
| 86% | 0.099 | 0.756 | 0.032 | 0.975 |
| 96% | 0.097 | 0.737 | 0.018 | 0.991 |

### Validation

The musculoskeletal simulations were validated by qualitatively comparing the simulations using the nominal model at 1.33 m/s, 3.0 m/s, and 4.0 m/s to the experimental data from Falisse et al. [1] (1.33 m/s) and Hamner & Delp [2] (3.0 and 4.0 m/s; Figure S 1, Figure S 2, Figure S 3). Additionally, cross-correlation coefficients and weighted root mean square errors (RMSE) were calculated between simulated and experimental time-series (Table S 7, Table S 8, Table S 9, Table S 10, Table S 11, Table S 12). RMSEs were weighted by experimental standard deviations (SD).

#### Walking - 1.33 m/s

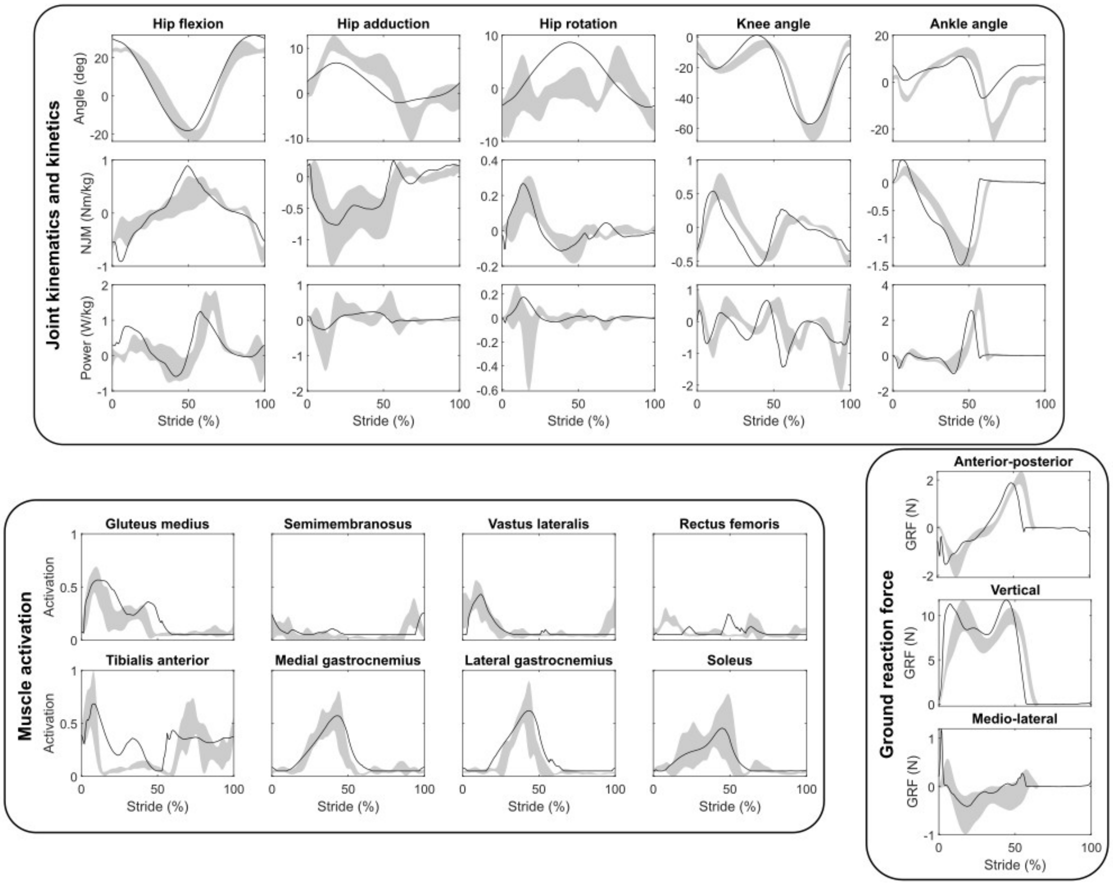


Figure S 1: Predicted time-series for hip, knee, and ankle joint angles, net joint moments (NJM), and powers, muscle activations, and ground reaction forces (GRF) during walking at 1.33 m/s using the nominal musculoskeletal model (black lines, compared to experimental time-series (mean ± 2 SD) from Falisse et al. [1]

Table S 7: Cross-correlation coefficients between simulated and experimental hip, knee, and ankle angles, torques, and power during walking at 1.33 m/s.

| Coordinate | Angle | Torque | Power |
|---|---|---|---|
| Hip - Flexion | 0.97 | 0.85 | 0.89 |
| Hip - Adduction | 0.94 | 0.96 | 0.67 |
| Hip - Internal rotation | 0.16 | 0.94 | 0.23 |
| Knee - Extension | 0.98 | 0.85 | 0.69 |
| Ankle - Dorsiflexion | 0.66 | 0.99 | 0.91 |

Table S 8: Weighted root mean square errors between simulated and experimental hip, knee, and ankle angles, torques, and power during walking at 1.33 m/s. The errors are expressed in units of standard deviations based on the experimental data simulation were validated against.

| Coordinate | Angle | Torque | Power |
|---|---|---|---|
| Hip - Flexion | 4.69 | 2.87 | 4.31 |
| Hip - Adduction | 3.27 | 2.31 | 2.05 |
| Hip - Internal rotation | 6.29 | 4.14 | 1.63 |
| Knee - Extension | 3.48 | 4.93 | 4.16 |
| Ankle - Dorsiflexion | 7.34 | 4.39 | 2.75 |

### Running - 3.0 and 4.0 m/s

Table S 9: Cross-correlation coefficients between simulated and experimental hip, knee, and ankle angles, torques, and power during running at 3.0 m/s.

| Coordinate | Angle | Torque | Power |
|---|---|---|---|
| Hip - Flexion | 0.96 | 0.90 | 0.82 |
| Hip - Adduction | 0.88 | 0.97 | 0.92 |
| Hip - Internal rotation | 0.59 | 0.92 | 0.74 |
| Knee - Extension | 0.97 | 0.85 | 0.77 |
| Ankle - Dorsiflexion | 0.95 | 0.99 | 0.96 |

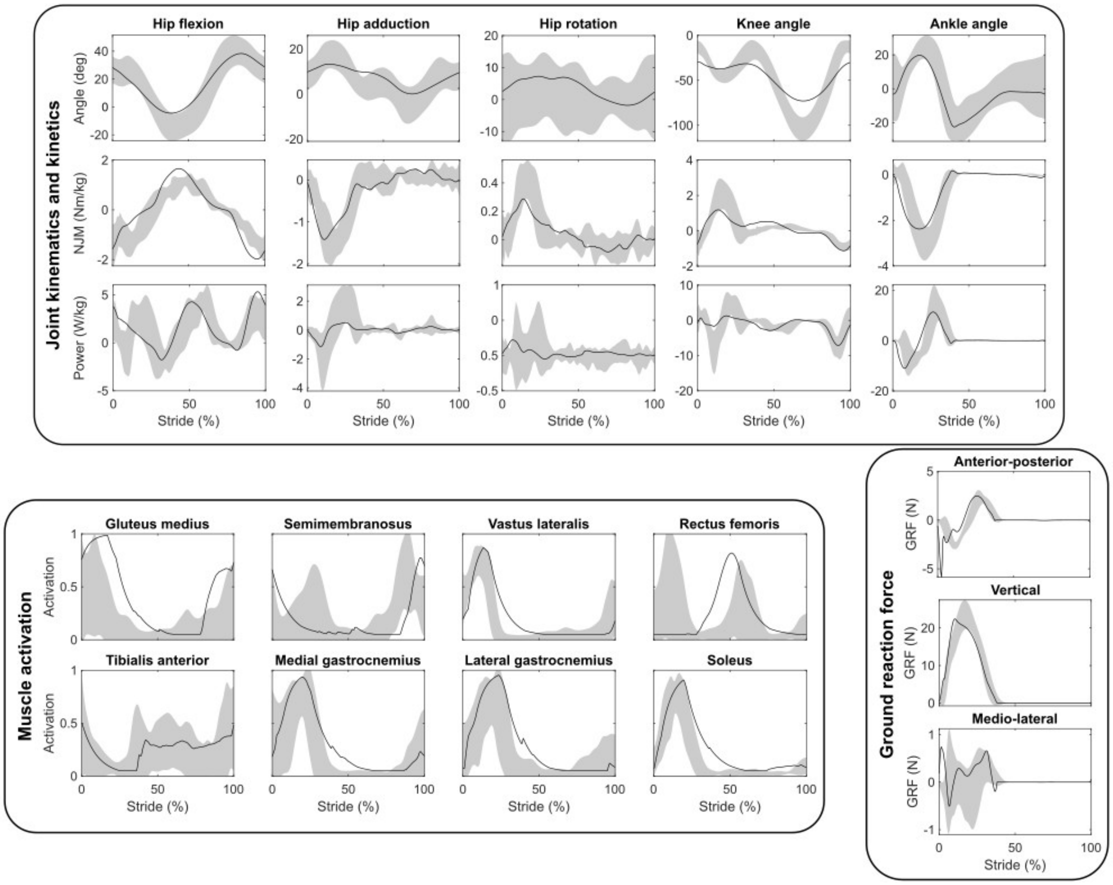


Figure S 2: Predicted time-series for hip, knee, and ankle joint angles, net joint moments (NJM), and powers, muscle activations, and ground reaction forces (GRF) during running at 3.0 m/s using the nominal musculoskeletal model (black lines, compared to experimental time-series (mean ± 2 SD) from Hamner & Delp [2]

Table S 10: Weighted root mean square errors between simulated and experimental hip, knee, and ankle angles, torques, and power during running at 3.0 m/s. The errors are expressed in units of standard deviations based on the experimental data simulation were validated against.

| Coordinate | Angle | Torque | Power |
|---|---|---|---|
| Hip - Flexion | 1.22 | 2.62 | 1.55 |
| Hip - Adduction | 1.13 | 1.36 | 0.85 |
| Hip - Internal rotation | 0.80 | 1.25 | 0.95 |
| Knee - Extension | 2.80 | 3.00 | 1.69 |
| Ankle - Dorsiflexion | 0.83 | 3.27 | 3.27 |

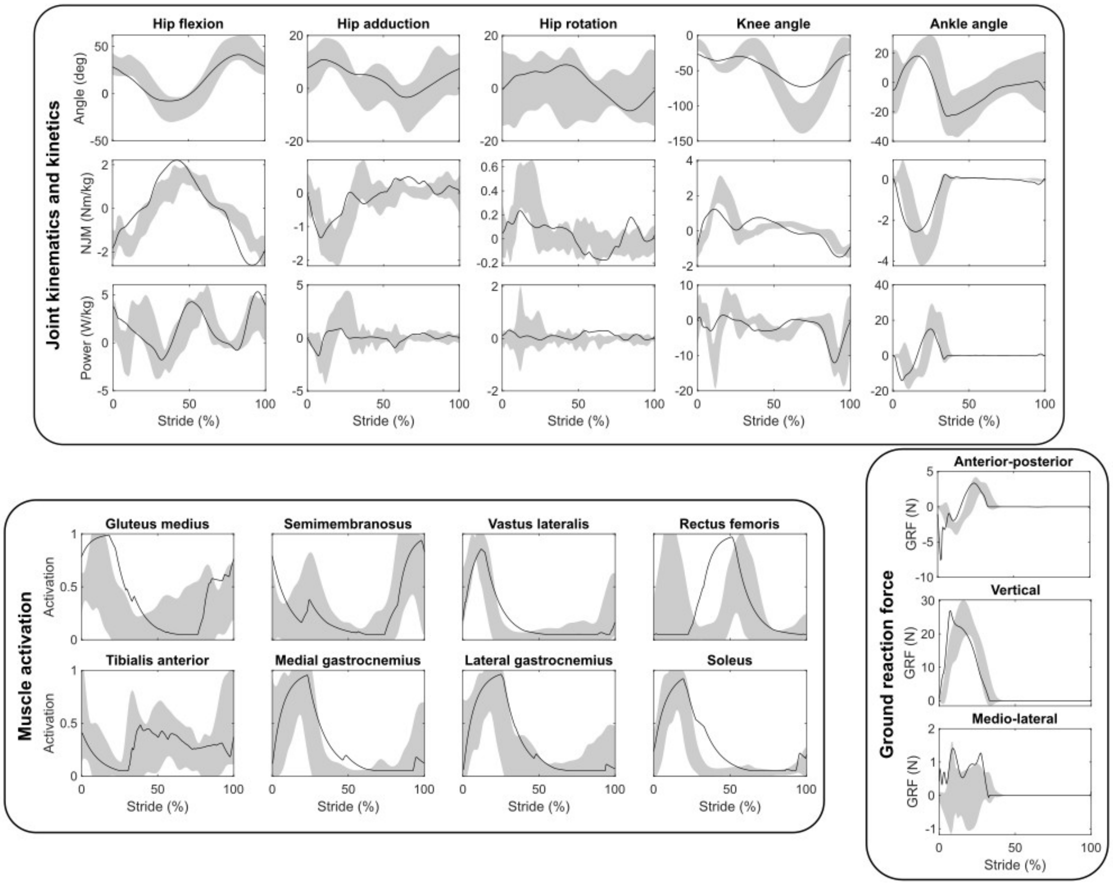


Figure S 3: Predicted time-series for hip, knee, and ankle joint angles, net joint moments (NJM), and powers, muscle activations, and ground reaction forces (GRF) during running at 4.0 m/s using the nominal musculoskeletal model (black lines, compared to experimental time-series (mean ± 2 SD) from Hamner & Delp [2]

Table S 11: Cross-correlation coefficients between simulated and experimental hip, knee, and ankle angles, torques, and power during running at 4.0 m/s.

| Coordinate | Angle | Torque | Power |
|---|---|---|---|
| Hip - Flexion | 0.97 | 0.87 | 0.84 |
| Hip - Adduction | 0.90 | 0.89 | 0.84 |
| Hip - Internal rotation | 0.59 | 0.63 | 0.35 |
| Knee - Extension | 0.97 | 0.79 | 0.85 |
| Ankle - Dorsiflexion | 0.95 | 0.99 | 0.96 |

Table S 12: Weighted root mean square errors between simulated and experimental hip, knee, and ankle angles, torques, and power during running at 4.0 m/s. The errors are expressed in units of standard deviations based on the experimental data simulation were validated against.

| Coordinate | Angle | Torque | Power |
|---|---|---|---|
| Hip - Flexion | 1.29 | 3.00 | 2.12 |
| Hip - Adduction | 0.85 | 2.21 | 1.35 |
| Hip - Internal rotation | 1.10 | 1.85 | 2.16 |
| Knee - Extension | 2.44 | 3.44 | 1.91 |
| Ankle - Dorsiflexion | 1.03 | 3.92 | 3.23 |

### Muscle fibre velocities

Muscle velocities were explored *post hoc* as a possible explanation of the different changes in thermodynamic efficiency between slow and fast twitch fibres (see manuscript for details). The muscle velocities of selected muscles important during locomotion are presented in Figure S 4.

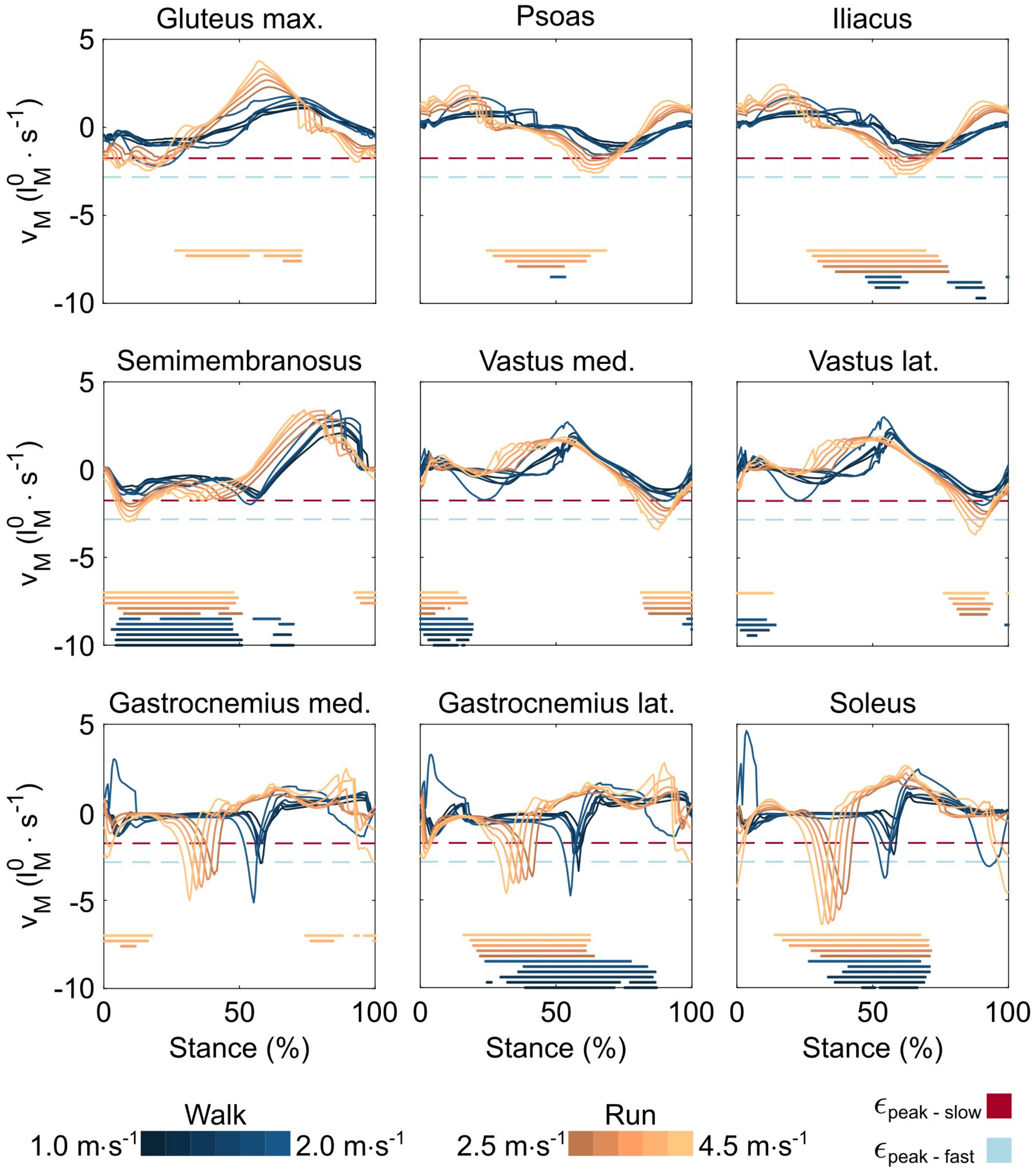


Figure S 4: The muscle shortening velocities of the nominal model's gluteus maximus, psoas, iliacus, semimembranosus, medial and lateral vastii, medial and lateral gastrocnemii, and soleus muscles during walking (blue curves) and running (orange curves). The lines underneath the plots indicate the times at which activation is greater than 0.5. Red and light blue dashed lines represent the shortening velocity at which peak efficiency for slow and fast twitch fibres occurs, respectively. $l_m^0$ is optimal muscle fibre length
